# Real-world visual search goes beyond eye movements: Active searchers select 3D scene viewpoints too

**DOI:** 10.1101/2025.02.08.637269

**Authors:** Tiffany C. Wu, John K. Tsotsos

## Abstract

Visual search is a ubiquitous task; people search for objects on a daily basis. However, the majority of the existing visual search literature focuses on passive search on a 2D computer screen, a far cry from emulating a real-world environment. Search is a real-world task that involves active observation. Search targets may be occluded, completely out of the observer’s line of sight, or oriented in unconventional ways. This is typically mitigated by actively selecting viewpoints, an important aspect of search behaviour with limited scope on a computer screen. Our goal was to explore viewpoint selection in active visual search. Subject eye and head movements were tracked as they moved freely while searching for toy objects in a controlled 3-dimensional environment, yielding the first such record of search-driven viewpoint selection. We found that subjects utilized their full range of eye and head motion to move from viewpoint to viewpoint, apparently employing a variety of objectives including changing viewing height and pose depending on object 3D pose. Subjects were also adept at selecting unobstructed views to search through otherwise occluded areas with objects. Furthermore, subjects completed the search task with high accuracy, even with no training on the environment. Although no learning was found in terms of accuracy over the duration of the experiment, increases in efficiency were found for other metrics such as response time, number of fixations, and distance travelled, particularly in target present trials where the target was not visible from the starting location. These results paint the story of a visual system that selects and moves to useful and informative views to facilitate the successful execution of an active visual search task, and stresses the significance of active vision research in understanding how vision is used in naturalistic environments.

## 1 Introduction

Although a large body of research exists studying human visual search, most of it has focused on the 2D, passive version of the task [1–4]. The classic visual search paradigm with 2-dimensional displays of artificial stimuli made by experimenters (popularized by Treisman & Gelade [2]) has since been used in many experiments (as seen in [1]), and is a mainstay of most psychology textbooks. This classic paradigm involves asking subjects to detect if a target stimulus is present or not, while measuring their reaction time and varying the number of distracting stimuli in the scene. Typically, in this task, the stimuli remain on the screen until the subject responds. Performance is measured by calculating the search slope (reaction time over set size), and is used to determine the difficulty of that search task. Paradigms where the stimulus onset period is manipulated are popular as well, and subjects’ accuracy is used to gauge the difficulty of the task.

While the ability to control for many factors is appealing, it is also a shortcoming of the search paradigm. Not only is depth limited to one image plane (unless it is simulated with various depth cues), the search space is also restricted to a confined area, meaning head or body movements may not be necessary, sometimes even prohibited or controlled, to perform search. In some experimental setups, even eye movements are not allowed.

Some of the contrast between this paradigm and the real world can be resolved using naturalistic images — images from the real world (as seen in [5–7]). As ’t Hart et al. [8] state, there is an issue with this approach — it is not known how informative static 2D images are for inferring gaze allocation in the real world. Tatler et al. [9] also argue that models based on such static picture-viewing paradigms do not encompass true gaze allocation behaviour in the real world, particularly when they model free-viewing tasks.

Research has only recently begun to focus on fully understanding active human visual search behaviour in a 3D environment [10–14]. Such an environment requires subjects not only to search, but to overcome issues of viewpoint selection, navigation, path planning, and more. Thus, in order to gain a better understanding of human visual search strategies in the real world, we must involve active vision and extend visual search to a 3D active observation task.

### 1.1 Active vision

Active vision is a key component of performing search in the real world. Bajcsy [15] defines active sensing as “the problem of intelligent control strategies applied to the data acquisition process which will depend on the current state of data interpretation and the goal or task of the process”. Some problems that are ill-posed and difficult to solve for passive observers become well-posed and easily solved with active vision [16]. Examples include shape from shading, shape from contour, shape from texture, and structure from motion.

Although not highly representative of real-world vision, tasks involving images of natural scenes can be active vision tasks as long as subjects can make eye movements. Indeed, many have sought to model eye movements during image search, including [17–19]. Any vision task involving a subject selecting views, whether by moving just their eyes, eyes and head, or even their body, is an active vision task.

Ballard [20] summarizes the computational advantages gained from “animate vision” versus passive vision. Any model of active vision, he asserts, must function in real time, being lightweight and highly dependent on adaptive behaviours. For a historical perspective, see Bajcsy et al. [21].

With this active vision perspective, we focus on human vision and eye and head movements [22–24]. The human head may exhibit 3 kinds of movement: dorsal or ventral flexion, tipping the head forward or backward about 60° in each direction; about 40° in left and right lateral bending or pivoting; and, left or right rotation, or panning, of about 65° either way [25]. These motions can also be described as combinations of movements along six dimensions of a head-centred coordinate system — translation in x, y, z (elevation) coordinates (partially due to head and partially due to body movement), and rotations *ϕ* (pitch — flexion), *θ* (roll — bending), *ψ* (yaw — panning). Note that there is no pure roll or pitch without some accompanying translational motion. Within the head, the eyes also move: abduction-adduction of about 55° (pitch — looking away or towards the nose); supraduction-infraduction of about 50° (yaw — looking up or down) [26]; and, up to 7° for incyclotorsion and -9° for excyclotorsion [27] (roll around the eye’s optical axis to focus above or below eye level).

One may assert that there is a canonical eye-head pose: with an origin at the center of the head pitch, yaw, and roll values of (0,0,0) and eye optical axes convergent at infinity in the horizontal plane through the eye centers, and zero cyclotorsion. Such a canonical pose is reasonable. Gravity provides ’vertical’ while the horizon is perpendicular to this. Balanced and minimal muscle tension sets the eye and head positions. Zero eye cyclotorsion is seen when the eyes fixate on points within Panum’s fusional area (horizontal and vertical), making it the best region for binocular fusion [24]. It may be suggested that much visual learning largely proceeds at such a canonical position and/or is intended to generalize to it.

This is the space of possibilities from which a human observer selects their next view of a scene. Our experimental work seeks to discover the range and characteristics of such selections while solving a 3D visual search task.

### 1.2 Active vision for navigation or exploration

One aspect of gaze allocation not captured in 2D viewing is gaze for navigation. The need to look in the direction of travel is not captured in 2D scenes, as no navigation is necessary. Significant research has been done on gaze control in the real world that focuses on navigation and eye-head coordination to direct gaze, but little for visual search [8, 10, 28–36]. Matthis et al. [28] found interesting results regarding gaze allocation and control in subjects walking on different types of terrain. In difficult terrains, fixations to where a subject is walking take up a significant proportion of fixations in the environment. This then influences gaze distribution in the real world in comparison to a 2D scene. Similar results have been found relating terrain complexity to gaze behaviours in [32] and [33].

Franchak et al. [10] further conducted a study investigating the difference in gaze distributions between walking and searching. Using IMU sensors as well as mobile eye trackers, they were able to determine the amount of head movement compared to eye movement during both tasks. They found that the search task induced a larger horizontal spread of eye and head movements. They also found subjects moved their head with significantly larger variability than their eyes during search.

Van Andel et al. [29] and Bonnet et al. [30] showed differences in gaze allocation when completing a task on a computer screen while standing versus sitting. Although these differences depended on the task, it is an indication that our gaze allocation in the real world while moving and active are likely different from sitting and facing a computer screen.

### 1.3 Active vision for specific goals

One of the first experiments investigating active search in humans was the classic study by Yarbus [37], where eye movements were tracked in subjects presented with an image while tasked with different queries regarding the image contents. He showed that eye movement patterns would differ drastically based on task demand.

Hayhoe & Ballard [38] provides a review of subsequent eye movement research in natural behaviours. They place emphasis on using active visually guided tasks to anchor meaning behind the fixations, rather than using only fixational data from experiments conducted on 2D screens (such as those seen in [37]). Active visually guided experiments can be conducted with the use of portable eye tracking devices, and studies have shown further that fixations are tightly coupled with task demand.

Recently, Solbach & Tsotsos [39] conducted an active same-different experiment to investigate active visual behaviours. They found that subjects select viewpoints in a way that suggests they are dynamically forming and testing hypotheses for the given task.

Ballard et al., Pelz et al., Hayhoe et al., and Land et al. [40–44]’s works combine both eye tracking, head movements, and actions. These experiments investigate eye, head, and hand coordination during a real world task. In [40] and [41] it was a block copying task, in [42], subjects had to make a sandwich, and in [43], subjects had to make tea. All three tasks involved search in the process, as subjects had to find the correct block to place, the knife or peanut butter jar when making the sandwich, or the sugar or milk for tea. One key insight from these experiments is that subjects acquire the information needed to perform each component of the task just prior to its use, rather than far ahead of time. They also found that the size of head movements is likely related to the constraints of the experiments, and that gaze shifts are almost always coupled with head movements in natural tasks. Land & Hayhoe’s review [44] further concludes that eye movements in these tasks are almost always to task-relevant locations, and are more driven by task demand than bottom-up salience. Indeed, Henderson [45] expresses in an opinion paper that viewpoint selection is driven by predictions based on knowledge or experience of the likely locations of task-relevant objects.

### 1.4 Active visual search

While eye and head movements during real-world navigation have been studied, there has been less research done in the same area focusing on visual search. Although Franchak et al. [10] compared eye and head movements when walking to search, they used a basic search and retrieval task, where potential targets were obvious 2-dimensional fabrics stuck on trees or benches. This meant that once the target was in the subject’s field of view, it would be easily identified without needing many more viewpoints.

There have been other visual search experiments conducted in a physical world setting [11, 12, 14, 46]. In Howard et al. [11], Sauter et al. [12], and Gajewski et al. [14], the search space was always a table or area directly in front of the subjects. In Foulsham et al. [46], the search space was a wall of mailboxes. In either case, there were very few, if any, occlusions that would require multiple viewpoints from a subject to verify a target’s presence and identity. In general, real world studies using active subjects requires specialized tracking and recording apparatus, and lacking these, many have approximated the real world using virtual reality.

### 1.5 Active visual search in virtual reality

Several active search experiments have been conducted in virtual reality [47–52]. Kit et al. [47] focused on scene structure guiding search and memory of objects in a more naturalistic environment, while Li et al. [48] took it a step further and compared the memory of objects with specific rooms in 2D versus an immersive reality environment. Others have also investigated the differences between virtual reality and the physical environment [52], as well as 2D natural scenes [50]. Olk et al. [49] also showed that a classic flanker study could be replicated with similar results in virtual reality compared to 2D.

Although they did not use a visual search task, Drewes et al. [53] showed comparable results between walking in virtual reality versus the real world in a path following task. However, head position and orientation was not tracked in the real-world task, and only eye movements were analyzed for comparison. Furthermore, the virtual reality headset only tracked head rotations (yaw, pitch, roll). There was no physical navigation through the world, only simulation via hand controller.

While a virtual reality environment offers many possibilities for investigating active observation, a real-world environment equipped with appropriate tracking capabilities, such as PESAO [54] or the STAR METHODS seen in [28], will provide higher ecological validity. Furthermore, there is no guarantee that subject eye and head movements in a virtual environment correspond to the same movements in the real world [55], even if performance measures may demonstrate some similarities [49, 52]. This is further shown in [56], where learning in navigation was found to be slower in virtual reality (head-mounted display with proprioceptive cues for walking and turning from an omnidirectional treadmill) compared to the real world, suggesting that differences still exist between the two types of environments.

### 1.6 Summary

Although these works are a step towards understanding active visual search in the real world, they have not addressed the importance of head and body movements to select viewpoints during search. In most experiments involving search, subjects are required to stay in the same location, and the search/task display is presented to them on a table, with only head and arm movements needed to complete the task. Several of the degrees of freedom in normal movement are limited or restricted in this design. If the butter knife was not on the experiment table in [42], where would the subject look, and how would they move to search for it if they could leave their seat?

Objects in real-world environments are frequently occluded, and a change in viewpoint by walking or moving the head is necessary to view all potential target locations and placements, particularly in a cluttered environment.

Thus, while much work has been done on search behaviour when the display is fully in view, how viewpoints are selected in order to locate and identify objects during search has not been well studied.

To address this gap, we conducted an active visual search experiment in a real-world environment. The PESAO experimental environment and apparatus [54] satisfied our real-world requirements (as it did for [39]), and thus approximations using VR or other means was not necessary. This environment is a controlled physical space, with synchronized mobile eye gaze and head tracking, enabling us to easily control the search space while allowing the freedom of movement for subjects to conduct active search. PESAO has been previously used in Solbach & Tsotsos [39].

Our primary goal with this experiment was to explore and measure human eye- and head-movement behaviours during active visual search in the real world. Specifically, we aimed to discover the characteristics of viewpoint selection during active visual search. Our secondary goal involved understanding different factors that may affect viewpoint selection, including experience with the search environment, object placement and orientation, as well as target presence, set size, and target visibility from the starting location.

## 2 Materials and methods

### 2.1 Experiment environment

The experiment was conducted in the PESAO [54] environment, a 3x4m space surrounded by blackout curtains, with 5 light sources and 6 Optitrack camera trackers. Multiple light sources were used to ensure more uniform lighting, and the Optitrack cameras were used to track the target location as well as the observer’s head movements at a rate of 120Hz. Observers were equipped with a set of Tobii Glasses 2 with Optitrack markers attached to track both their eye gaze and head movements during the task. Subjects were untethered and free to move as they wished in the space. The total head mount weight was 66.6g (glasses are 45g, Optitrack markers mount is 21.6g) [57], and the recording unit subjects had to place in their pocket or clip to their clothes was 312g. Target objects were tagged with another custom set of Optitrack markers, and their location was recorded before the start of each trial, and then the markers would be removed to avoid markers being used to cue target location. Tobii Pro records the first person view from the glasses during each trial, and an additional camera was set up to record the entire scene during the experiment, such that another vantage point would be available for matching recorded measurements with physical actions during manual analysis. In other words, both the first person view and third person views of each trial were captured on video.

The Tobii Glasses 2 were calibrated once at the start of the experiment. Gaze event data was directly exported from Tobii Pro Lab, including fixations, saccades, their respective durations, as well as their 2D and inferred 3D coordinates from the eye images. The eye trackers had a sampling rate of 50 Hz, and gaps in the eye gaze data up to 75ms were interpolated. A fixation was defined as a sequence of raw gaze points with estimated velocity below 30°/s. Fixations and saccades were defined on an eye-in-head basis. We manually labelled fixations that landed on specific objects, as well as the target object. A fixation was labelled as “object” or “target” as long as the fixation point as plotted in Tobii Pro Lab landed anywhere on the object. The Tobii Glasses 2 have an average precision error of 1.29° while walking [58].

According to [59], the Tobii Glasses 2 at 50 Hz were minimally affected by slippage of the tracker on subjects’ faces, making them a good candidate for use in a task involving subjects walking around and moving their head. Furthermore, the measurements we are taking from the eye gaze data include fixation location and duration, as well as saccade amplitudes across whole saccades, all two-point temporal samples whose sampling errors are centred around 0, according to [60]. Additionally, our focus is more on where people are looking, rather than specific timings of particular gaze events. As such, the precision afforded by a 50Hz tracker is sufficient for the purposes of our analyses in this experiment.

Head location and orientation (tracked using custom markers attached to Tobii Glasses), as well as target location (tagged with custom Optitrack markers), was directly exported from Motive Optitrack, giving the xyz location as well as orientation (roll, pitch, yaw) of the rigid body, in the head-in-world basis. The Optitrack system has a measurement error of *±*0.2*mm*.

For every subject, the data streams from the Tobii Glasses 2 and Motive Optitrack were saved, as was a controller script with each trial’s start and end times. Using their respective timestamps, the three data streams were merged, with missing values interpolated across time using the custom PESAO software (see [54] for more detail). After the merging, a single large dataset including all eye gaze and head movement data, along with markers for the start and end of each trial, was generated. The code and data for this experiment is available at https://osf.io/vqdb2/.

Targets and distractors were palced within cages and on the table surfaces. Tables and wireframe cages were arranged in pre-determined configurations in the area, such that observers would have to walk and navigate around to obtain different viewpoints during the search task.

Wireframe cages were chosen so that the Optitrack cameras would still be able to record markers for target objects through the cage walls, rather than being occluded by solid cage walls. Black coverings were placed on some faces of the cages, increasing the amount of occlusion to encourage subject movement through the setup, while ensuring that enough cameras could see into each cage to maintain accurate tracking. The tables and wireframe cage locations stayed the same throughout the experiment for each subject, but they varied between 4 different versions across subjects (Fig 1).

**Fig 1.**
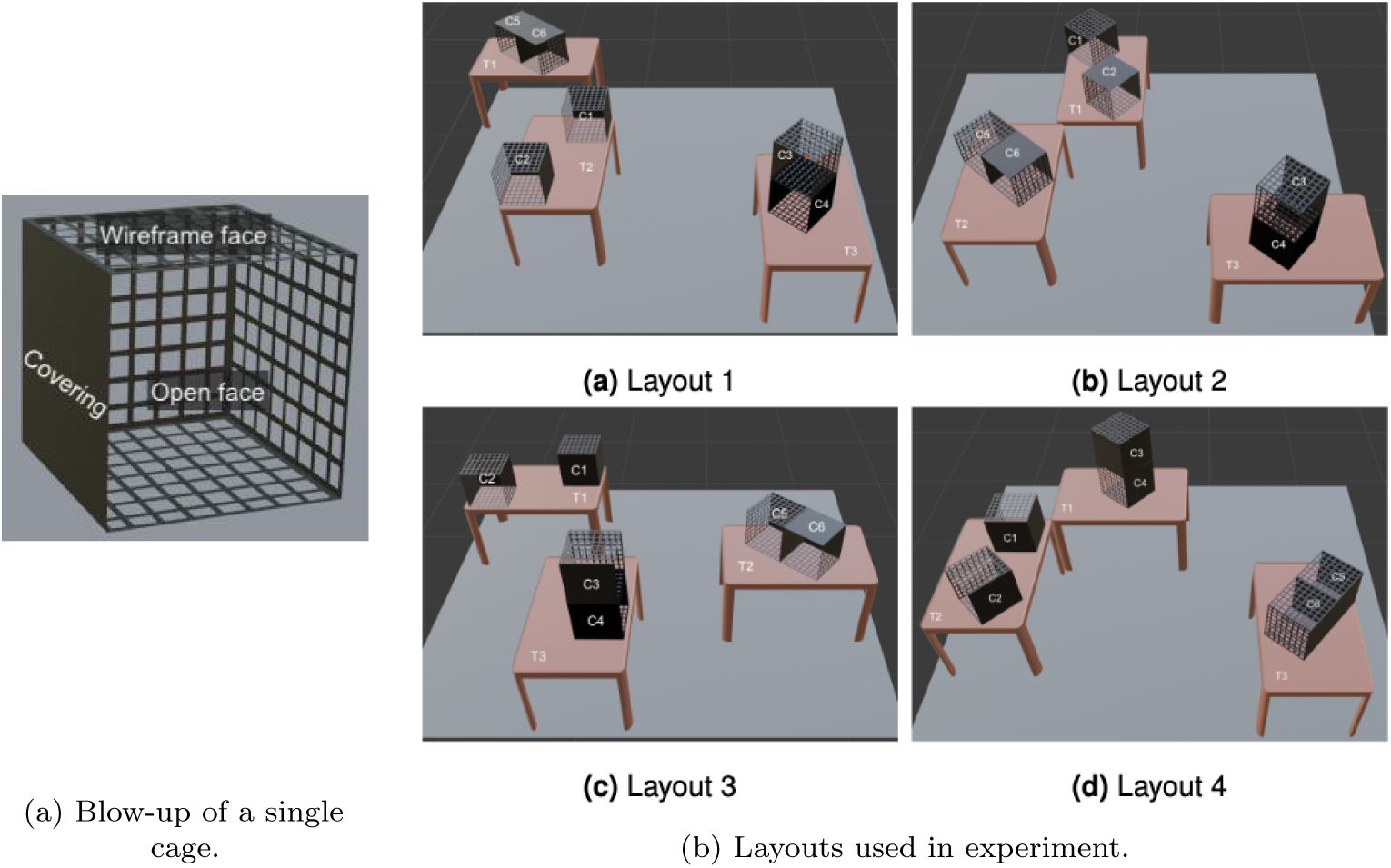
Layouts used in the experiment. Each table and cage has a label, and objects are assigned to a particular label in each trial. The blow-up in (a) shows that solid sides are covered faces, sides with the grid texture are wireframe sides, and there is one open face with no cover at all. Cage dimensions are 30x30x30cm, while tables are 114(w)x70(d)x70(h)cm.

The layouts of the tables and the cages were not chosen randomly. There are a vast number of potential configurations of tables, cages, and objects. This particular sampling was chosen in order to make the task tractable, with the assumption that it is representative of the configuration space. The number of tables and cages was kept constant across the layouts to make for easier analysis and trial generation. 3 tables were chosen to allow for enough space in between the tables, regardless of the many configurations possible, for people to walk through the setup. 6 cage shelves were chosen as that was the number that gave sufficient table space and cage space with the 3 tables. Each setup included a tower of 2 cages, with the second cage being stacked above the first, to increase the vertical height variance of the search space. Each setup also included a unit of 2 cages horizontally connected, as well as 2 cages that stayed as individual cages.

As in all common visual search experiments, our subjects were asked to determine if a specified target is present or not. The target was introduced to the subject as an image on a monitor at the start of each trial for 5 seconds while the subject was facing away from the search environment. All targets were presented in their canonical view. This view was determined by the experimenter, and showcased the objects in poses that seemed representative and made the objects distinguishable. Once the image disappeared, subjects could turn around to begin the search. Subjects could move freely within the experimental setup in order to complete the search (see Figure 2), and there was no time limit.

**Fig 2.**
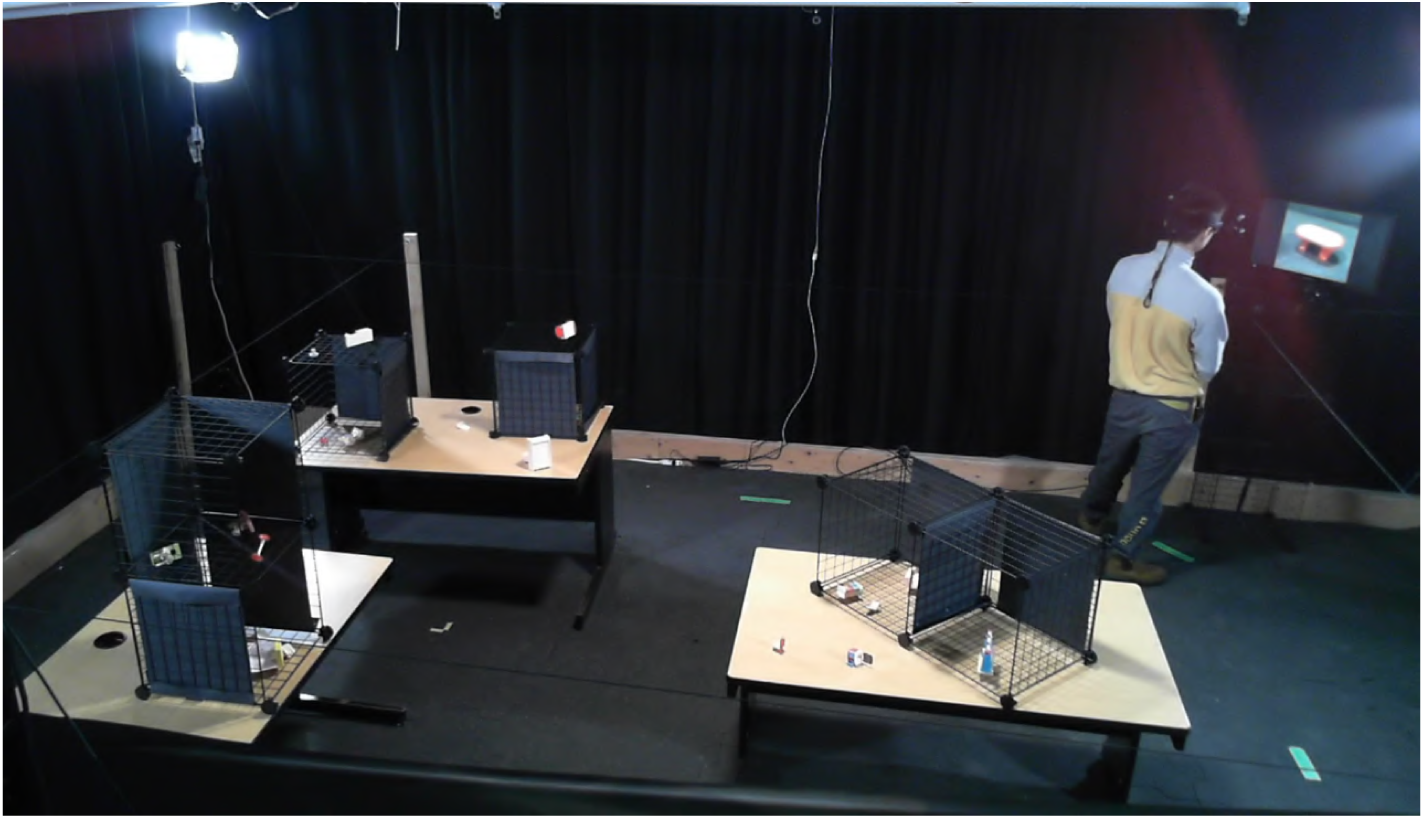
Experiment environment example — layout 3. The subject is viewing the target on the monitor. This display lasts for 5 seconds. The subject then turns around and searches for the target.

Once the subject decided, they had to verbalize their response. If they believed the target was present, they were told to fixate on the target, while saying “I have found the target”. If they believed the target was absent, they had to say “I think the target is absent”. Subjects began from the same location for every trial (in the far right corner — top right in plots in Figures 2, 1), and each subject performed 12 trials in total.

### 2.2 Ethics statement

93 subjects (21 excluded) participated in the experiment. The experiment was approved by the York University Office of Research (STU 2023-004). All subjects gave written informed consent. All subjects had normal or corrected-to-normal vision and were not colour-blind. Subjects were recruited from January 23, 2023 until November 18, 2024.

### 2.3 Experiment design

A 4x6 between-subjects design was used to counterbalance the table-cage layout and trial version for the 72 valid subjects. There were four table-cage layouts and six trial versions, yielding 24 combinations of the two factors. Three subjects were run on each combination. The number of valid subjects needed was determined a priori to ensure enough data was collected to analyze appropriately.

Stimuli were 1:12 size dollhouse furniture of everyday objects, such as chairs, tables, couches, etc (see Fig 3 for some examples). They are made in a variety of colours, materials, and sizes. These stimuli are not qualitatively different from blocks or kitchen items like those used in past active vision experiments [41–43], and were chosen in order to achieve a large set size of unique looking objects in the scene. Images of the stimuli were taken in a canonical view such that they would be easily recognizable when presented to subjects as their targets. Objects ranged from 1.5cm to 15.1cm in length, width, and depth.

**Fig 3.**
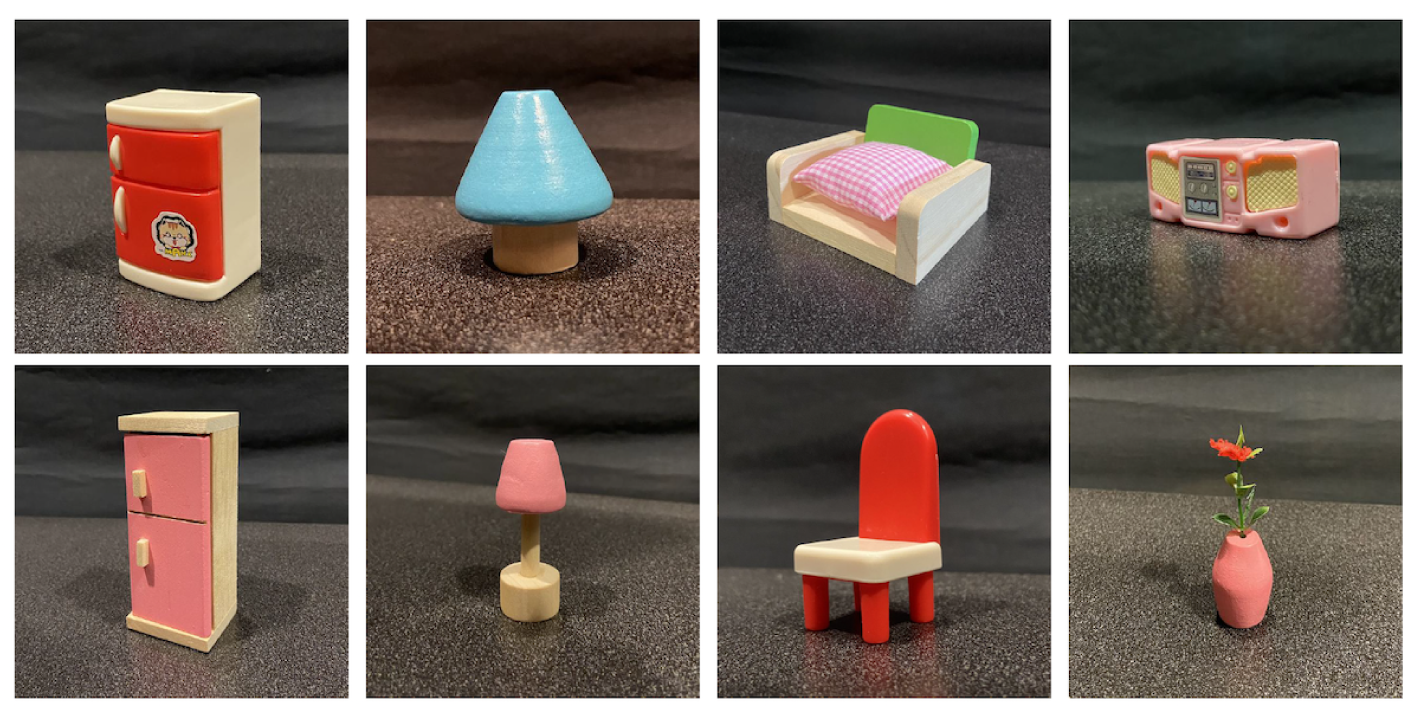
Example of some stimuli used in the experiment.

Stimuli were placed in pre-determined locations in the setup for every trial, and no trial had the same set of objects in the same locations. The orientation of all objects was randomized, while the orientation of target-present objects were chosen a priori and stayed constant across subjects. There were 6 different sets of 12 trials, and subjects were run on one of the six. Targets and distractors were randomly chosen from a pool of 119 objects. Although these toy objects are artificial, we chose to use them in order to have a higher level of experimental control. As scene grammar has potential to guide search [61], and we were using furniture objects, we made sure to disrupt potential scene grammar cues by varying object orientations and randomizing the types of objects that would coincide in the same table or cage. We also took careful note of the objects’ colour, shape, size, and material in order to balance out the distribution of objects in each trial.

Three independent variables were manipulated:

1. Target presence [present, absent]
2. Target visibility from starting location [yes, no]
3. Set size [30, 40, 50, 60]

To further elaborate the target visibility variable, this was defined as whether or not the target object was visible when standing at the marked starting location, when the subject is facing towards the tables and cages after they turn around.

Of these variables, we fully counterbalanced target presence within the 12 trials for each subject, and counterbalanced the visibility and set sizes between subjects, such that there were always an even distribution of set sizes and target visibilities, but they were paired to different trial orders across the different trial versions. We then generated objects for each trial given the set size, and whether the target was present. After generating the objects for the first trial, they were randomly assigned a table or cage (chosen from 9 possibilities: table 1, table 2, table 3, cage 1, cage 2, cage 3, cage 4, cage 5, cage 6) such that there were approximately the same number of objects in each (we allowed a tolerance of *±*2). Subsequent trials were generated such that as many objects were shared between consecutive trials, given the trial order. This was done to minimize the time subjects needed to wait between trials as we changed the stimuli in the setup. If a target was present, it would always be removed in the trial immediately after. Objects new to the next trial are then assigned a location in order to balance out the number of items in each location as much as possible. There were never 2 trials with the same target object in each trial version. Code used to generate trial versions can be found here: https://gitlab.nvision.eecs.yorku.ca/tiffwu/active-search-trials-generator.git.

We measured subjects’ response time, accuracy, eye gaze data, and head movement data. We also recorded the location of the tables and cages, as well as target objects.

## 3 Results

93 subjects participated in the active visual search experiment. 4 were excluded due to too many invalid gaze samples from the Tobii eye tracker (*<*65% valid across both eyes, where average validity was 85.8%, *σ* = 8.7%), 10 were excluded due to missing data points transmitted from the Motive Optitrack head tracking system, and 7 were excluded due to interrupted Tobii eye tracking recordings. Of the included subjects, 32 were male, and 40 were female.

There were four different layouts of tables and cages used across subjects. In each layout, there were 3 tables and 6 wireframe cages in which objects were placed in random 3D poses. Fig 2 shows one such layout. Objects changed in between each trial such that subjects would not be able to memorize the locations of previously seen objects.

Subjects completed trials in time ranging from 4 seconds to 3 minutes long (there was no time limit), and there were a total of 864 trials combined across subjects. In target-present trials, subjects took 50.5 fixations, travelled 9.3m, and took 24.6s on average. In target-absent trials, they took 102.8 fixations, travelled 19.4m, and 47.5s on average. The total experiment, including the time it took to arrange objects between trials, ranged from 20 minutes and 54 seconds to 44 minutes and 34 seconds.

In the following analyses, we calculated the 3D view vectors of each fixation by combining head and gaze direction and logged at which of these 9 locations (tables or cages) each fixation was directed at, or if they were not directed at any. Fixations directed to tables or cages are termed “look-at” fixations. The head pose used for each fixation was averaged over all the samples taken during the fixation, as was the eye gaze direction. Target fixations were manually labelled, as mentioned earlier.

Aside from the raw fixation, saccade, and subject head location and orientation data, the following metrics were also used in analysis, and were all extracted from head location and orientation data from the Motive Optitrack system:

**Crouches** Times when a subject’s recorded height (elevation of tracking apparatus) is less than 2SD below their average height across the experiment

**Head tilts** Subject head roll and pitch angles

**Distance travelled in 3D (m)** Total distance a subject travels across 1 trial

**Revisits** Number of revisits a subject takes to a table or cage in the setup

We visualized subjects’ search-paths in 3D, as shown in Figs 4 and 5. A search-path is the subject’s head trajectory (the dashed white line), with head pose for each fixation over the course of a trial. Each fixation is represented by a viewing frustum, with the opening (wide end) indicating the direction of gaze. See the blow-up of the frustum in the bottom-left corner of Fig 4 for clarification. The pointed end of the frustum is one endpoint of the eye optical axis; the black line shows gaze direction pointing out from this apex. Grey frusta are fixations looking at the environment, blue frusta are fixations looking at a table or cage, and red frusta are target fixations. The path of the subject is represented with the dotted white line that links the recorded head poses. The first and last frusta are labelled “start” and “end”. The target, if present, is indicated with a bright red dot, labelled “target”. Videos of subjects performing the experiment from both a third person view and from a first person view are available here: https://osf.io/vqdb2/.

**Fig 4.**
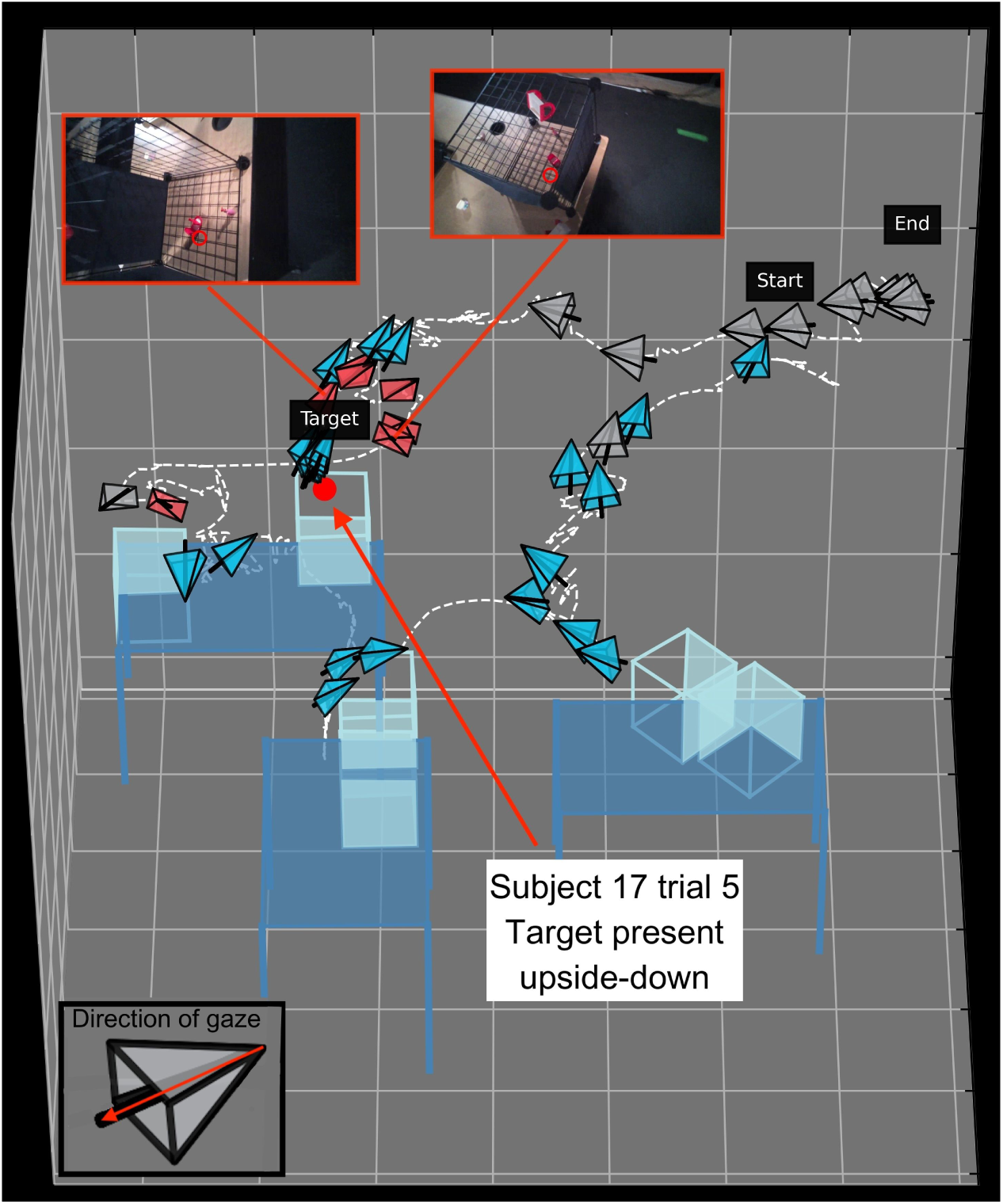
Scanpath of subject 17 performing a trial. Darker blue rectangles and lines indicate the tables, and lighter blue cubes indicate cages. Opaque faces of cubes are occluded faces. Each fixation is represented by a viewing frustum with the opening indicating direction of gaze. The blow-up of the frustum in the bottom-left corner has a red arrow overlaid to show how to interpret direction of gaze from the frustum. Environment fixations are grey, table or cage look-at fixations are blue, and target fixations are red. The target was present (shown by red circle), and it was placed in the environment upside-down. Once the subject spotted the potential target, they oriented their view to confirm. Images embedded in the figure are enlargements of the particular fixation image they are connected to, with the red circle within indicating their fixation point. These show two different views from the subject fixating on the target before their response.

**Fig 5.**
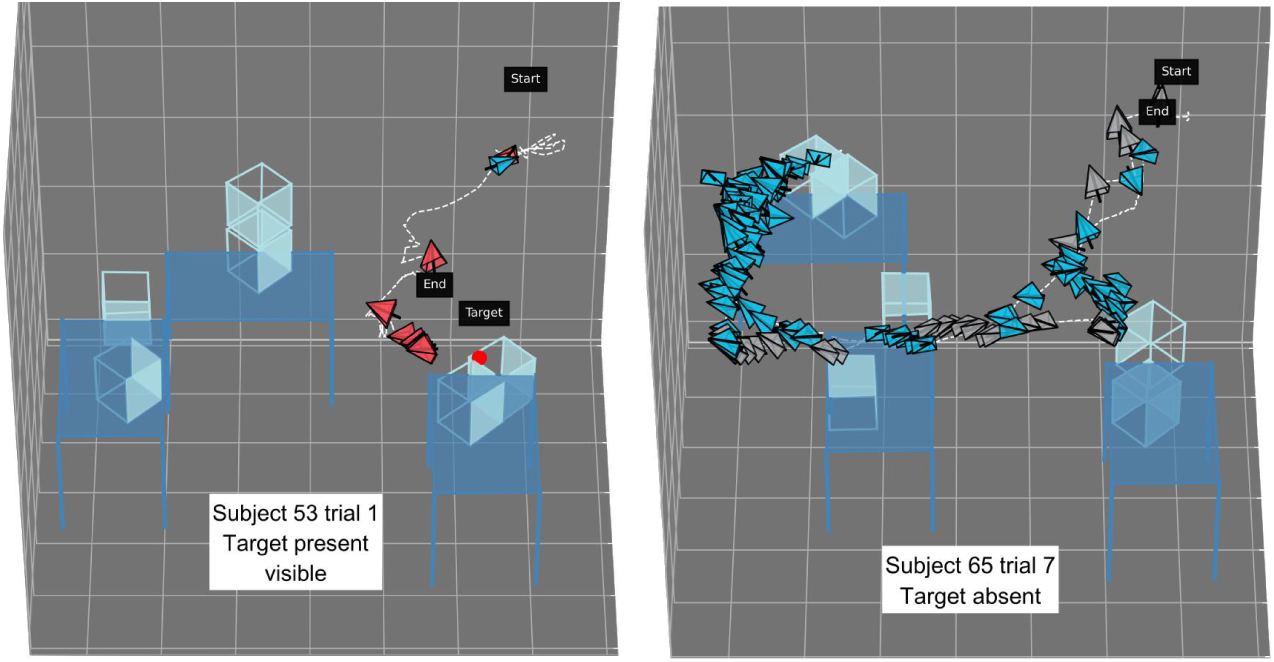
Scanpaths of a target present (visible) and target absent trial. In the target visible trial from subject 53, the subject immediately discovers the target, approaches, and confirms its presence, then terminating the search without exploring the rest of the environment. In the target absent trial from subject 65, the subject walks through the search space to explore the environment with many fixations before determining the target is absent.

### 3.1 Effect of layout and trial version

4 layouts and 6 trial versions were chosen to mitigate the influence of a specific placement or trial on overall results. To ensure these layouts and trials were sufficiently different, we conducted a 2-way ANOVA (layout x trial version) on number of fixations, response time, and distance travelled. All three metrics were log-transformed so as to correct to a normal distribution. Significant interactions were found for number of fixations (*F*_15,840_ = 1.79*, p* = 0.032, partial *η*^2^ = 0.031) as well as distance travelled (*F*_15,840_ = 1.75*, p* = 0.037, partial *η*^2^ = 0.031). The only significant pairwise difference found through Tukey’s HSD was between (trial version 3, layout 4) and (trial version 6, layout 2) for number of fixations.

In terms of response time, there was a significant main effect of trial version (*F*_5,840_ = 3.46*, p* = 0.004, *η*^2^ = 0.019), and the only significant pairwise difference was between trial version 1 and 6. A significant main effect for layout was also found (*F*_3,840_ = 4.22*, p* = 0.006, *η*^2^ = 0.014). Here, there were no significant pairwise differences.

Thus, to test the effects of table, cage, and object configurations, we performed ANOVAs showing the significant effects of layout (table and cage configurations), as well as trial version (object configurations). Results reported in the following sections are aggregated across layout and trial versions, and characteristics of active search behaviour were similar across these configurations, leading us to believe these characteristics are rather generalizable. However, it is important to keep in mind that any discovered characteristics may not extrapolate to edge cases, such as environments with only one object, or with no occlusions whatsoever.

### 3.2 Eye and head movement characteristics in active visual search

Subjects take a new view of the environment with each fixation. To gain a basic understanding of subjects’ viewpoint selections, we first present analysis on the number of fixations, general statistics about the fixations and saccades, as well as the types of views chosen.

When the target was present and subjects responded correctly, the number of target fixations was 4.2 on average (*σ* = 3.89). The mean number of consecutive target fixations immediately prior to making their response was 2.45 (*σ* = 1.94).

Average fixation durations were 199.6ms (*σ* = 165), compared to the typical 200-330ms found in 2D search experiments, including scene image search [62, 63]. When comparing to other real-world experiments [36, 42, 43], we noticed that several did not report fixation durations. For those that did, duration averages had a wide range, from 215ms to 420ms. However, characteristics such as large spread and long right tails were consistent with what we found.

After removing outliers (durations *>*3 SD above or below the average), we also found target fixation durations (*µ* = 295.75*ms, σ* = 242.5) to be significantly longer than other fixations’ durations (*µ* = 176.4*ms, σ* = 102.8), with a two samples t-test yielding *t*(64226) = 29.1*, p <* 0.000.

With the combination of eye gaze and head tracking, we were able to compute saccade amplitudes with both eye-in-head and head-in-world data. Firstly, eye-in-head saccade amplitudes were on average 10.4 ° (*σ* = 8.03), which is higher than in 2D natural image search, where the average saccade amplitudes are around 5-6° [63].

Eye-in-world saccade amplitudes, computed by adding the eye-in-head to the head-in-world data, gave average amplitudes of 46.2° (*σ* = 55.3). This is a larger amplitude and spread than reported in [10], although their search targets were placed in a much larger search space and allowed detection from further away. Thus, this could simply be a result of differences in the design of the experiment environment.

Breaking down the eye-in-world (eye + head) saccade amplitudes, we see that the head contributes more than the eyes in 52.9% of all instances. The median proportion of eye contribution per saccade is about 46.6%. Saccades with no head movement or minimal head rotation (less than 1°) take up 15.2% of all saccades, comparable to the 19% reported in [35] during real-world exploration. Fig 6 shows the distribution of saccade amplitudes from the eye and the head separately after removing head amplitudes of 0. We can see clearly that eye amplitudes taper off by 50°, while head amplitudes have an extremely long tail going all the way to 175°. This range of eye vs. head amplitudes are consistent with prior literature [31], which state that humans have an oculomotor range of about 55° to each side.

**Fig 6.**
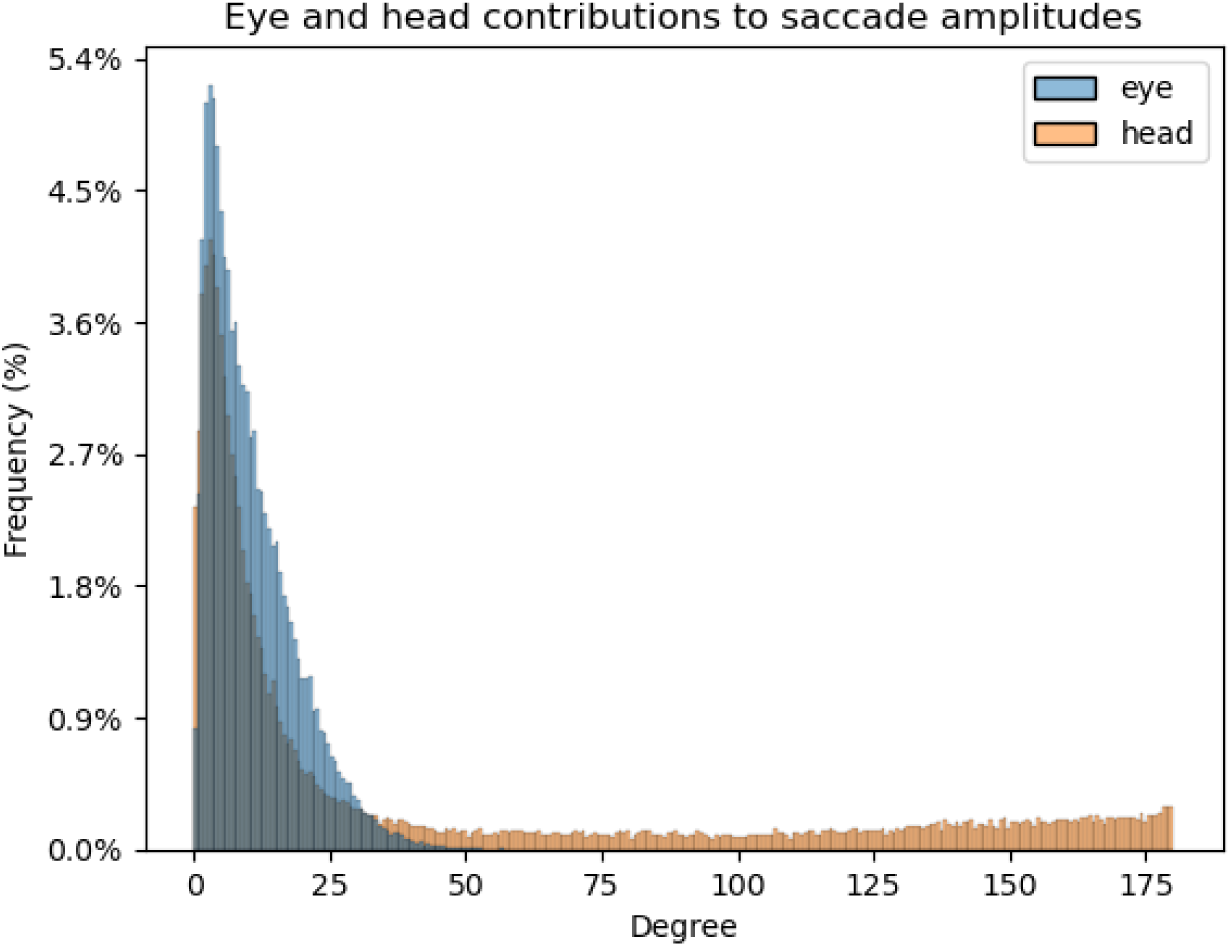
Distribution of eye and head saccade amplitudes overlaid. Eye amplitudes are concentrated within 50 °, while head amplitudes have a long tail to 175 °

The wide range of eye and head amplitude distributions suggest that real-world searchers are utilizing the full degrees of freedom of their eyes and head. With 2D search on screens, or even search on a table in front of the subject, the useful range of eye gaze and head movements are smaller, and thus may not be representative of subjects’ true search behaviours in the real world. Indeed, a typical 24-inch computer screen subtends only 56 ° when a subject is seated 57cm away. Unfortunately, other studies performing visual search in virtual reality have not reported subject eye and head amplitudes, so it is unknown if they would be comparable.

Regarding the direction of gaze (eye-in-world), we can divide fixations into “look-at” fixations, where the subject is looking at a table or cage (where all the stimuli are), and environment fixations, where the subject is not looking at any tables or cages. 75% of fixations are “look-at” fixations, and 25% are environment fixations. Indeed, these environment fixations are not captured in 2D search — any understanding of real-world search must combine how fixations for searching and fixations for navigation are decided and integrated into an overall search strategy.

We then analysed the distribution of head rotation in the form of roll, pitch, and yaw values across trials. Rolls are left and right lateral bending of the head (although Optitrack only captures the rotation). Pitches are dorsal or ventral flexion — tipping the head forward or backward. Yaws are turning the head left and right.

After removing outliers (angles more than 3 SD from mean, typically occurring due to noise from Optitrack marker recordings), average roll angles measured over the course of the experiment for all subjects were 0.7 ° (*σ* = 23.4) and average pitch angles were -5.3 ° (*σ* = 33). This is on par with results found measuring head orientation statistics during everyday activities in [64], with a lower spread for roll angles as well as a negative average pitch angle, indicating subjects are tilting their head slightly down. This is consistent with the search environment, as tables and cages were situated below eye level. The spread of both roll and pitch angles are larger in this dataset compared to [64], and this may be due to the amount of head movement induced by the search environment compared to the everyday activities recorded in [64]. Average yaw angles were 6.3 ° (*σ* = 95.1), as subjects had free range to look in all directions in the environment. Fig 7 shows the three distributions, and Fig 8 shows roll and pitch distributions on a polar plot.

**Fig 7.**
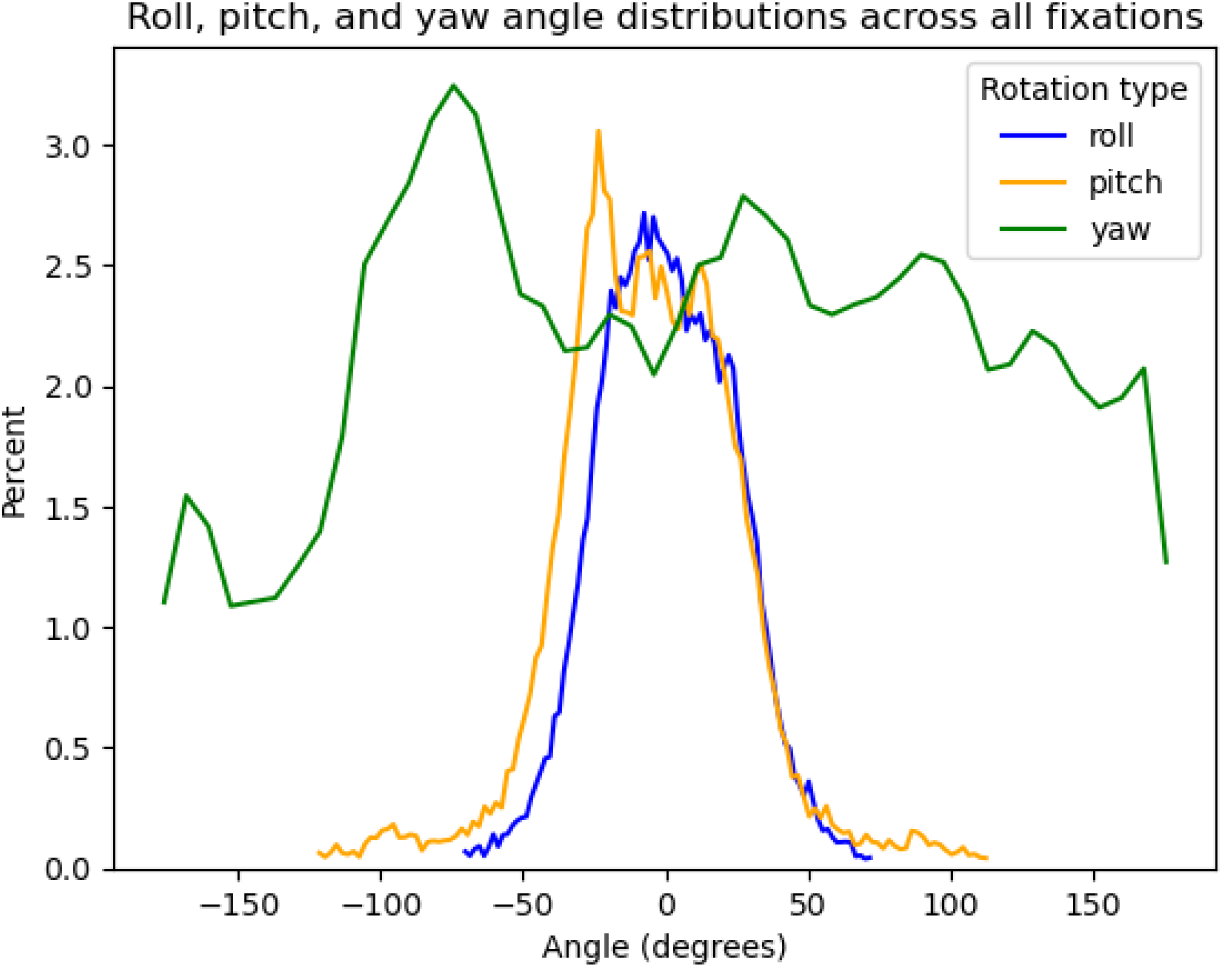
Distribution of roll, pitch, and yaw angles across all fixations from all subjects.

**Fig 8.**
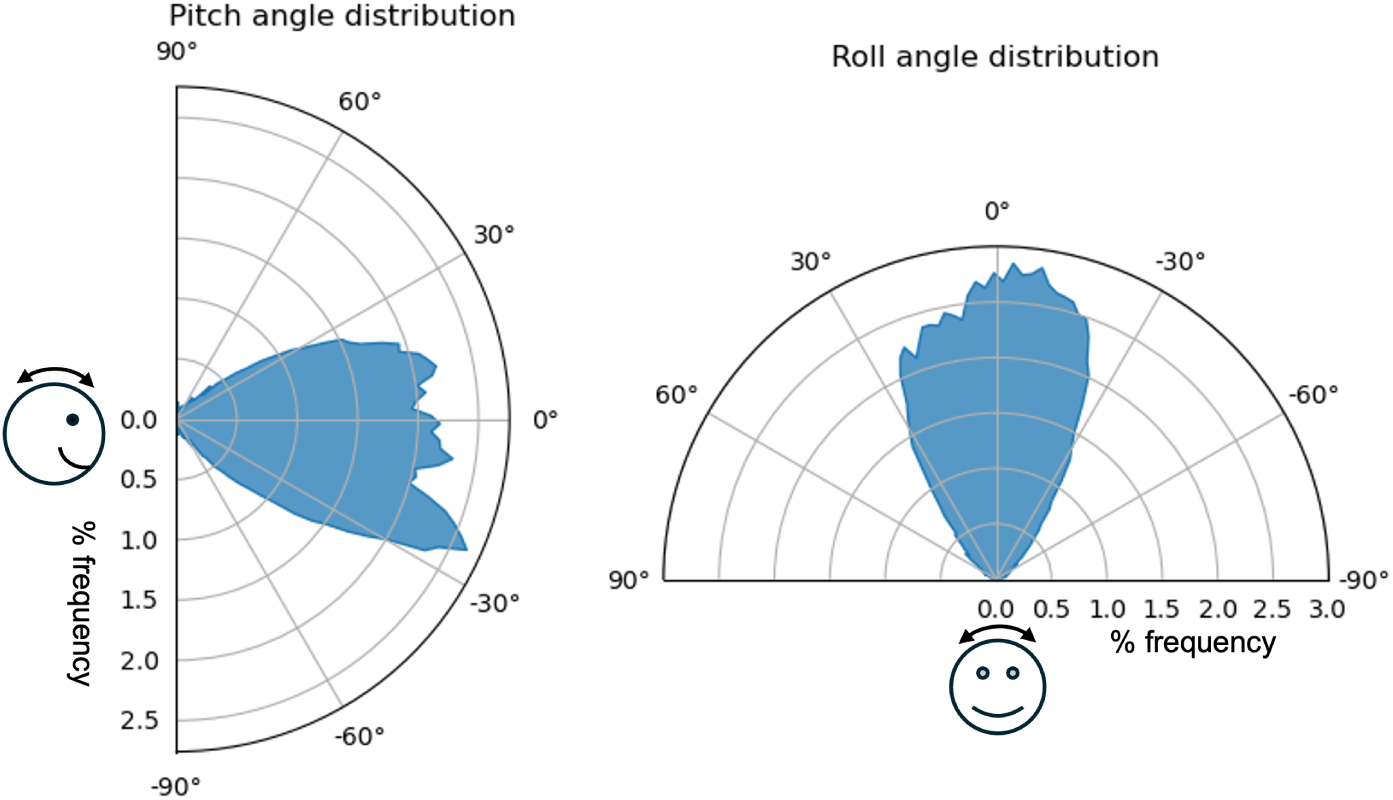
Roll and pitch distributions on a polar plot. Roll distributions are relatively even in both directions (*µ* = 0.7°), whilst pitch angles are on average more negative (*µ* = *−*5.3°), reflecting the tendency for subjects to tilt their heads slightly down to view the tables and cages in the search environment.

As mentioned in Section 1.1, the typical human head and neck range of motion is around 40° in lateral bending, and 60° in forward or backward tilting. Thus, the range seen in Fig 7 for rolls and pitches corroborates that subjects use their full range of head tilting motion during the search task.

As we logged the exact table or cage at which each fixation was directed, we can further analyze revisits, where a subject views a table or cage, leaves to view a different table or cage, and eventually returns. The number of revisits to a table or cage in target-present and absent trials were 5.3 (*σ* = 6.3) and 18.2 (*σ* = 13.8) respectively.

Considering target-absent trials, the average number of revisits to each specific table or cage across different layouts varied from 0.41 to 4.89, while the average number of unique location revisits was 5.94 (out of 9).

Although subjects tend to perform several revisits to tables and cages, they do not fixate on every object in the scene directly, even in target-absent trials. There are 134 out of 432 target-absent trials where the number of fixations is fewer than the number of objects in the scene. When manually labelling look-at fixations, we noticed that although subjects looked in most of the areas with objects, they may skip some objects in a cage or a table. It is possible that subjects fixate in a general area, then decide that they do not need to perform any further inspection there because they did not see any objects of the right colour or shape to match the target. It is also possible that the information subjects gained from their peripheral vision indicated that there were no hints of target presence, and there were no candidate targets that triggered fixations.

#### 3.2.1 Summary

We documented several characteristics of viewpoint selection in active visual search. Fixation durations were slightly shorter than those found in other real-world experiments, although the distribution and spread were similar. We also found longer fixation durations towards targets compared to other locations. Gaze amplitudes were larger and had larger spread compared to [10], although this may be due to the design of the experiment environment.

The large spreads in eye and head amplitudes suggest that subjects use their full range of motion to conduct visual search and navigate the environment.

Finally, we show that subjects tend to make several revisits during their search, particularly during target-absent trials, even though they may not fixate on every object in the environment. This observation is novel, as previous real-world experiments involving visual search have not included target-absent trials.

These descriptive characteristics of eye gaze and head movement behaviours in active visual search contribute to the growing evidence that visual tasks in the real-world involve many more behaviours than those recorded on 2D versions of the same task.

### 3.3 Search accuracy and efficiency

Overall, subjects performed very well during the task (overall accuracy *µ* = 90.9%, *σ* = 8.8%), though their false negative rates were higher than their false positive rates (12.5% vs. 5.8% respectively). To determine whether experience affects viewpoint selection, we compared several metrics for performance and efficiency (response time, number of total fixations, number of non-environment fixations, distance travelled) over the course of the 12 trials, to see if any learning effect can be found.

No learning effect was found over the duration of the experiment. Subjects were just as accurate in their first trial as they were in the last (see Fig 9 for accuracy by trial number).

**Fig 9.**
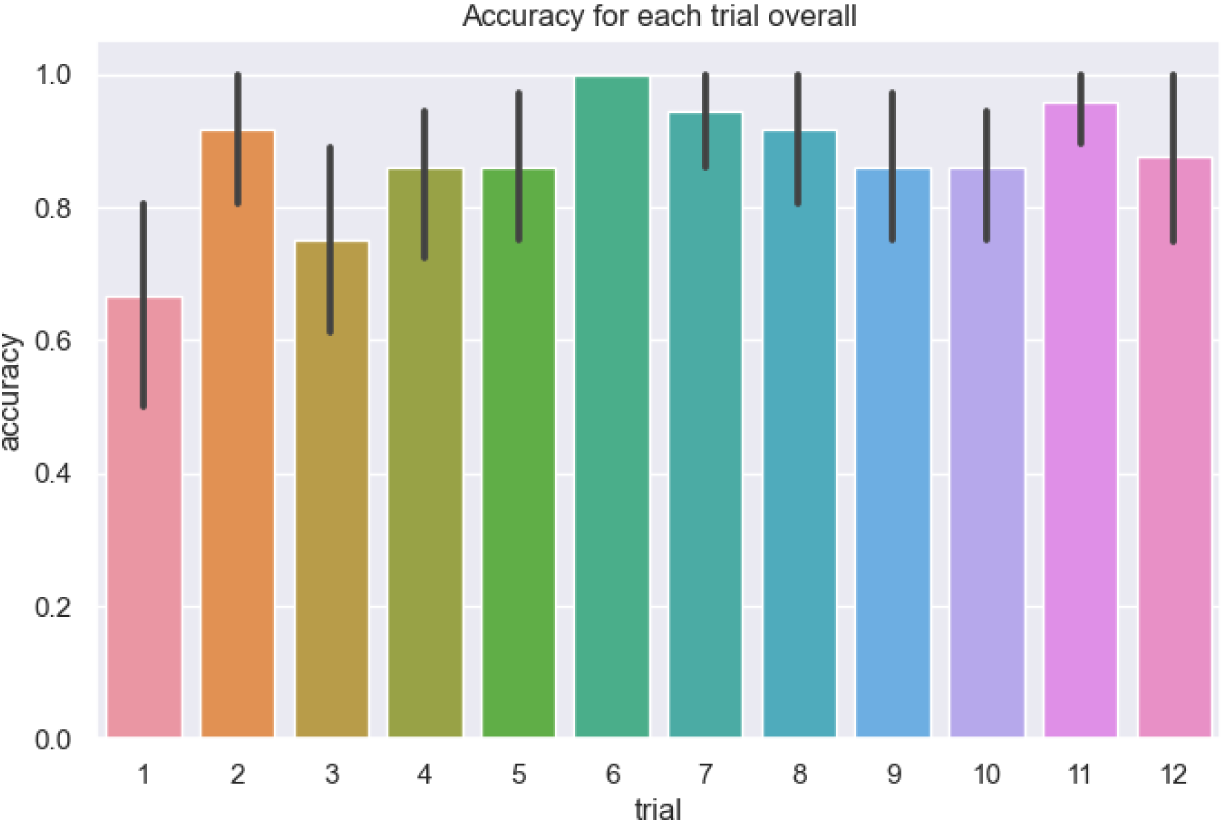
Average accuracy of each subject over trial number. Overall accuracy is very high, and there is no significant increase in accuracy over the trials. Black lines represent the 95% confidence intervals.

A significant increase in efficiency was found in terms of distance travelled during target-present trials, with a Spearman’s rank correlation coefficient of *r*(10) = *−*0.12(*p* = 0.017) between trial number and average distance travelled, number of look-at fixations (*r*(10) = *−*0.13*, p* = 0.012), number of total fixations (*r*(10) = *−*0.11*, p* = 0.03), and response time (*r*(10) = *−*0.1*, p* = 0.049). Although significant, the magnitude of these relationships is quite small. The negative relationships point to subjects becoming more efficient as the experiment went on.

There was no significant increase in efficiency in any measures for target-absent trials. Computed Spearman’s rank correlation coefficients between trial number and each measure are as follows: response time *r*(10) = *−*0.046*, p* = 0.35, number of fixations *r*(10) = *−*0.002*, p* = 0.97, number of look-at fixations *r*(10) = 0.03*, p* = 0.61, and distance travelled *r*(10) = *−*0.005*, p* = 0.91.

Target visibility was determined as whether the target was visible or not when viewed from the starting location, in the corner of the environment. When we further split the trials by target visibility, we also see no increase in efficiency for visible trials, with correlation coefficient magnitudes below 0.05 and p values above 0.5. However, for trials where the target was not visible, significant correlations between trial number and our efficiency metrics were found. For distance travelled, *r*(10) = *−*0.19*, p* = 0.008, for number of total fixations, *r*(10) = *−*0.17*, p* = 0.02, for number of look-at fixations, *r*(10) = *−*0.22*, p* = 0.003, and for response time, *r*(10) = *−*0.17*, p* = 0.02. Indeed, correlations were the strongest here, thus it seems that subjects do become more efficient in completing target not visible trials over time.

#### 3.3.1 Summary

To address an aspect of our secondary goal, determining whether viewpoint selection is affected by experience, there is some evidence for learning in terms of response time, number of fixations, and distance travelled that was observed only in target-present trials, perhaps because accuracy is near ceiling from the start. It seems that subjects are incredibly proficient at the task of searching in this unfamiliar environment even without any training at all. This can be compared to the result in Solbach & Tsotsos [39] where for a different visual task, they too found no learning effect in accuracy, but improvements over trials in response time, fixations, and head movement.

The significant relationships between trial number and all efficiency metrics for target not visible trials, but not for target visible trials, could be due to the larger margin for improvements possible in not visible trials, and the possibly quick termination of target visible trials if subjects spot the target from the starting location.

### 3.4 Effect of target presence, set size, and visibility

To determine the effects of target presence, set size, and target visibility on viewpoint selection, we quantified and broke down various eye gaze and head movements from the raw data as mentioned above, then conducted two 2-way ANOVAs to analyze their effects on these metrics. In particular, we consider the following metrics: trial response time, number of fixations, distance travelled (m), trial accuracy, and the number of revisits.

First, isolating target-present trials to include the target visibility variable, we conducted a 2-way ANOVA (target visibility x set size) on the metrics mentioned. Significant interactions were found for distance travelled (*F*_3,424_ = 3.05*, p* = 0.028, partial *η*^2^ = 0.021), accuracy (*F*_3,424_ = 3.21*, p* = 0.023, partial *η*^2^ = 0.022), and number of revisits (*F*_3,424_ = 3.42*, p* = 0.017, partial *η*^2^ = 0.024). Tukey’s HSD revealed several significant pairwise differences. The pair (30, visible) vs (60, visible) was significantly different both for distance travelled and number of revisits, with the (30, visible) trials having less distance and fewer revisits. (30, visible) vs (50, not visible) was also significantly different for distance travelled. For number of revisits, (30, not visible) and (60, visible) was significantly different. Visible trials at set size 60 had more revisits on average (*µ* = 11.2) than all other cases (4.5 *< µ <* 8.3). Finally, one pairwise difference was found between not visible and visible trials at set size 50 for accuracy, with visible trials having higher average accuracy (97% vs 78%).

Thus, it seems like for distance travelled, trials with set size 30 and visible target had less travelling compared to trials with larger set size (50 and 60). For revisits, trials with set size 60 and visible targets had more revisits than set size 30 trials. Finally, at the set size 50 level, it seems like visibility did affect subject accuracy.

For metrics with no significant interaction, there was a significant main effect of set size (response time *F*_3,424_ = 6.03*, p <* 0.001, partial *η*^2^ = 0.04, number of fixations *F*_3,424_ = 9.04*, p <* 0.001, partial *η*^2^ = 0.06). Tukey’s HSD revealed significant pairwise differences between set size 30 vs 60 as well as 40 vs 60 for both metrics.

Interestingly, there was no significant main effect of visibility from the starting location for these metrics, especially since trial efficiency differed by visibility in the earlier section. We then conducted further analysis on subject head locations in their first few fixations, and found that they move out of the starting location (alway far right corner of the space) quickly, after an average of 2.47 fixations (*σ* = 3.33). Moving out of the starting location was defined with a threshold of 30cm from the head location of their first fixation in each trial. Thus, many subjects left the position where the target was visible before fixating on the target, possibly leading to the lack of significant effect for visibility.

Contrary to our belief that subjects would observe the scene in some detail before forming a strategy for walking and moving, we see that they instead begin moving around to search almost immediately. This indicates a search strategy that uses active vision first, rather than one that uses the full range of eye and head motion only after a stationary scanning of the scene does not reveal a target object.

Secondly, we conducted a 2-way ANOVA (target presence x set size) on the same select metrics, this time using all trials. No significant interactions were found. Overall, there was a strong significant main effect of target presence (response time *F*_1,71_ = 141.9*, p <* 0.001, number of fixations *F*_1,71_ = 139.3*, p <* 0.001, distance travelled *F*_1,71_ = 185.6*, p <* 0.001, accuracy *F*_1,71_ = 6.85*, p* = 0.01, number of revisits *F*_1,71_ = 90.7*, p <* 0.001). Calculating effect size (*η*^2^) gave large values of around 0.26 for response time and number of fixations, with a smaller value of 0.21 for number of revisits. *η*^2^ was 0.35 for distance travelled, but only around 0.01 for accuracy.

A significant main effect of set size was also found for response time (*F*_3,213_ = 12.7*, p <* 0.001, *η*^2^ = 0.029), number of fixations (*F*_3,213_ = 22.4*, p <* 0.001, *η*^2^ = 0.049), distance travelled (*F*_3,213_ = 6.98*, p <* 0.001, *η*^2^ = 0.016), and number of revisits (*F*_3,213_ = 19.9*, p <* 0.001, *η*^2^ = 0.04). Tukey’s HSD revealed that for all these metrics, there was a significant pairwise difference between set sizes 30 and 60. For number of fixations and number of revisits, there was also a significant difference between set sizes 40 and 60. Finally, a significant difference between set sizes 50 and 60 were found for number of fixations, and a significant difference between set sizes 30 and 50 were found for number of revisits.

#### 3.4.1 Summary

Thus, to address another aspect of our secondary goal of establishing whether target presence, set size, and target visibility affect viewpoint selection, we discover that visibility interacts with set size to affect accuracy, distance travelled, and number of revisits, while target presence has the largest effect, and set size has some effect only for certain pairwise differences. The weaker significant effects of set size may be due to the presence of occlusions in the scene, meaning subjects needed a base amount of movement to search all the possible locations regardless of set size.

In terms of visibility from the starting location, subjects spend very few fixations (2.47) on average scanning before beginning to move around in the space, thus mitigating the measured effect of visibility. However, it is interesting that subjects are not always scanning for the target fully before moving. One possible reason for this behaviour is that objects ranged from 1.5-15.1cm in size, which, from the starting location, could range from 0.23 to 5.2° visual angle, depending on its size and where it is placed in the scene. As some objects may appear very small from the starting location, subjects may prefer to begin moving to get closer to those objects to perform a more effective search. Thus, getting closer to the objects becomes a more useful strategy to get the objects within a range where object recognition and search is easier. If the targets were highly salient and could pop out with a single glance, then perhaps always scanning at the starting location would be a better strategy.

### 3.5 Effect of object placement and orientation on viewpoint selection

To address the effect of object placement and orientation on viewpoint selection, we discuss the manipulation of viewpoints achieved by varying target orientations, as well as occlusions caused by covering different cage faces in different layouts.

#### 3.5.1 Target orientation

To further probe eye gaze and head movements during search, target orientation in 3D was varied. The target object’s image presentation always depicted a canonical view of the object. Orientations of targets in the scene included upright (matching canonical orientation as presented in image), front-up, side-up, diagonal, and upside-down (see Fig. 10). Shepard & Metzler [65] found a linear relationship between response time and depth-plane orientation differences of two objects during a same-different task, concluding that subjects could mentally rotate objects at an average rate of 60° per second.

**Fig 10.**
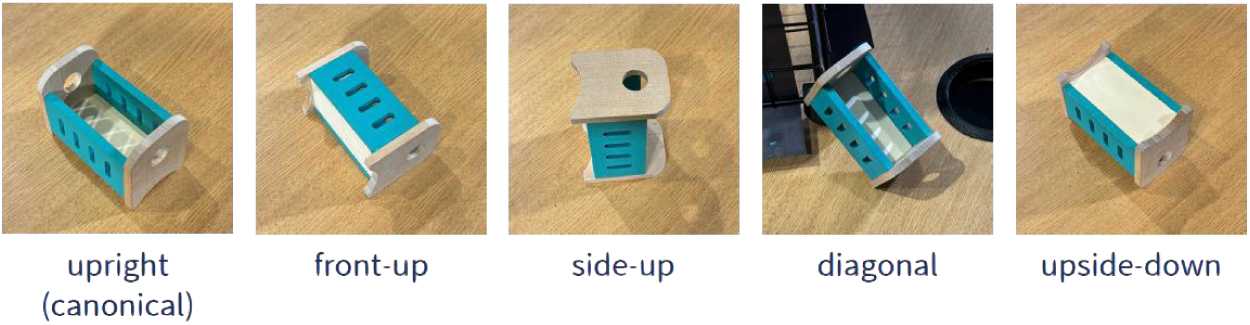
The five different orientations that target objects were placed in. The orientation of an object is dependent on the canonical image taken of the image. For objects with a distinct front and side, the front was always the left face in the canonical image. Objects with no distinct front or side, but were placed on their sides, were labelled front-up.

We hypothesized that, given the freedom to move the head to select viewpoints, subjects would orient their view to match an object to the target presentation during search, using both head tilts (roll and pitch) and elevation (whether a subject is crouching or not). From a standing position, diagonally oriented objects matched closest to the target image presentation. With regard to target height, targets could be placed on tables (0.7m from ground), on the bottom of a cage (same height as tables), on top of one cage (around 1.1m from ground), and on top of a stack of two cages (around 1.4m from ground).

One-way ANOVAs were performed on roll and pitch angles, as well as subject head elevation in target-present trials to test for the effect of target orientation. We first removed outliers (defined as *>*2SD from the mean angle or elevation), then performed three one-way ANOVAs.

A one-way ANOVA performed on the average roll angles in each target orientation revealed a significant difference between groups (*F*_4,1415_ = 2.47*, p* = 0.04, *η*^2^ = 0.007). No significant pairwise differences were found.

A one-way ANOVA was similarly performed on the average pitch angles by target orientation, revealing a significant difference between groups (*F*_4,1411_ = 3.43*, p* = 0.008, *η*^2^ = 0.01). For pitch angles, significant pairwise differences were found between side and upright orientations, with average pitch angles of -9.14 (*σ* = 44.9) for side and -1.55 (*σ* = 28) for upright. Negative values indicate subjects’ heads are tilted down.

Finally, a one-way ANOVA on subjects’ head elevation for each target fixation relative to their standing height (estimated head height when standing straight) over all trials revealed a statistically significant difference between different orientations (*F*_4,1401_ = 2.774*, p* = 0.026, *η*^2^ = 0.008). The standing head height was computed using the maximum elevation of the subject’s head across the experiment. Significant pairwise differences were found between the front (*µ* = 0.9*cm, σ* = 9) and side (*µ* = 2.5*cm, σ* = 6.48) orientation.

With regard to the placement of target objects, we performed a linear regression between subject head elevation (relative to their estimated standing head height) and target elevation to determine if subject head elevation would correlate to the elevation of the surface the target is placed on. A significant positive correlation (*r* = 0.343*, p <* 0.001) was found, with a slope of 0.76 and intercept of 0.95 (see Fig 11). Targets placed at table height (around 0.7m) corresponded with lower head positions than targets placed on top of the highest cage (around 1.4m).

**Fig 11.**
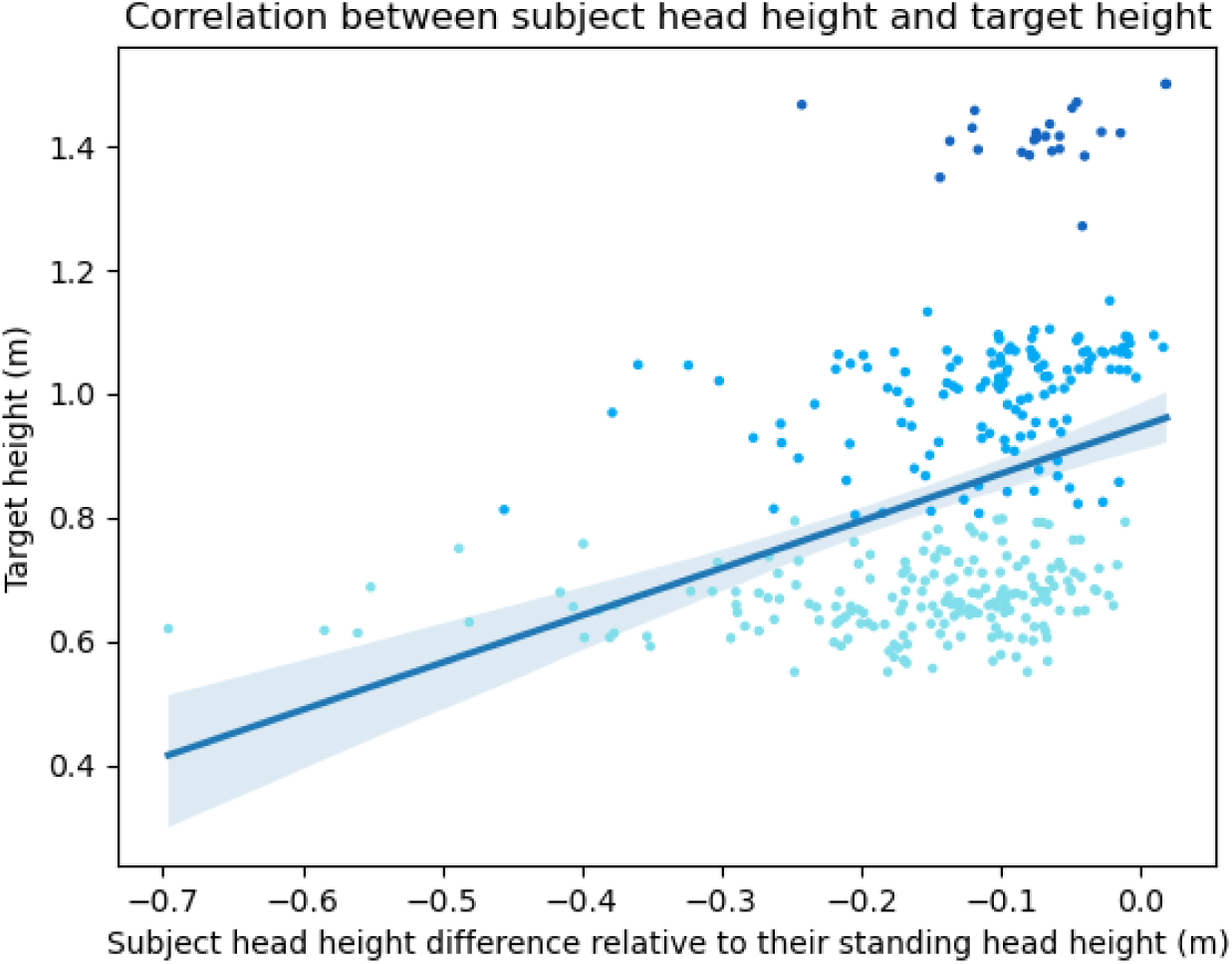
Correlation between subject head height and target height. Target heights look like they have steps because targets can only be placed on a surface. This leaves three possible target heights: table height (turquoise dots), table + one cage (light blue dots), and table + 2 cages (dark blue dots). Pearson’s correlation coefficient was *r* = 0.34 and significant, indicating that subjects’ head heights were lower when objects were placed lower, and vice versa.

As Blanz et al. [66] showed, the elevation of preferred object viewpoints in most cases is significantly below 45 °, meaning subjects rarely rotated objects to view them from above. This is reflected in the lowering of head elevation for objects placed lower in the scene, showing that subjects are selecting more preferable views during their search.

It is possible that there is not only a canonical view for objects, but also a canonical viewing geometry that is preferred by subjects. That is, if the human canonical viewing geometry is the one where the head has zero pitch, roll and yaw, and the optical axes of the eyes intersect the object-of-interest straight ahead (with equal vergence angle) and in the horizontal plane, then our subjects display a tendency to achieve this with their head movements.

#### 3.5.2 Effect of occluding faces

On each cage, certain faces were occluded using black coverings (see Figs 2, 1), causing objects to be visible only from certain viewpoints. These faces were chosen to allow for the manipulation of target visibility from the starting location, as well as to mimic real-world conditions where many target objects are occluded.

To determine if subjects’ fixation distributions were affected by visibility into cages, DBScan was used [67] to find clusters of subject head locations(epsilon=0.15, min samples = 2%), indicating locations that subjects spend more time standing to conduct their search. A hotspot was defined as a cluster from DBSCAN. Fig 12 shows each hotspot color-coded with their corresponding look-at location frequencies in the bar chart beside it. Hotspot locations in each layout correspond to the openings of cages. Each hotspot also has its peak fixation look-ats matching the cage openings that the hotspot is facing, indicating that subjects find views with the least amount of occlusion to conduct their search.

**Fig 12.**
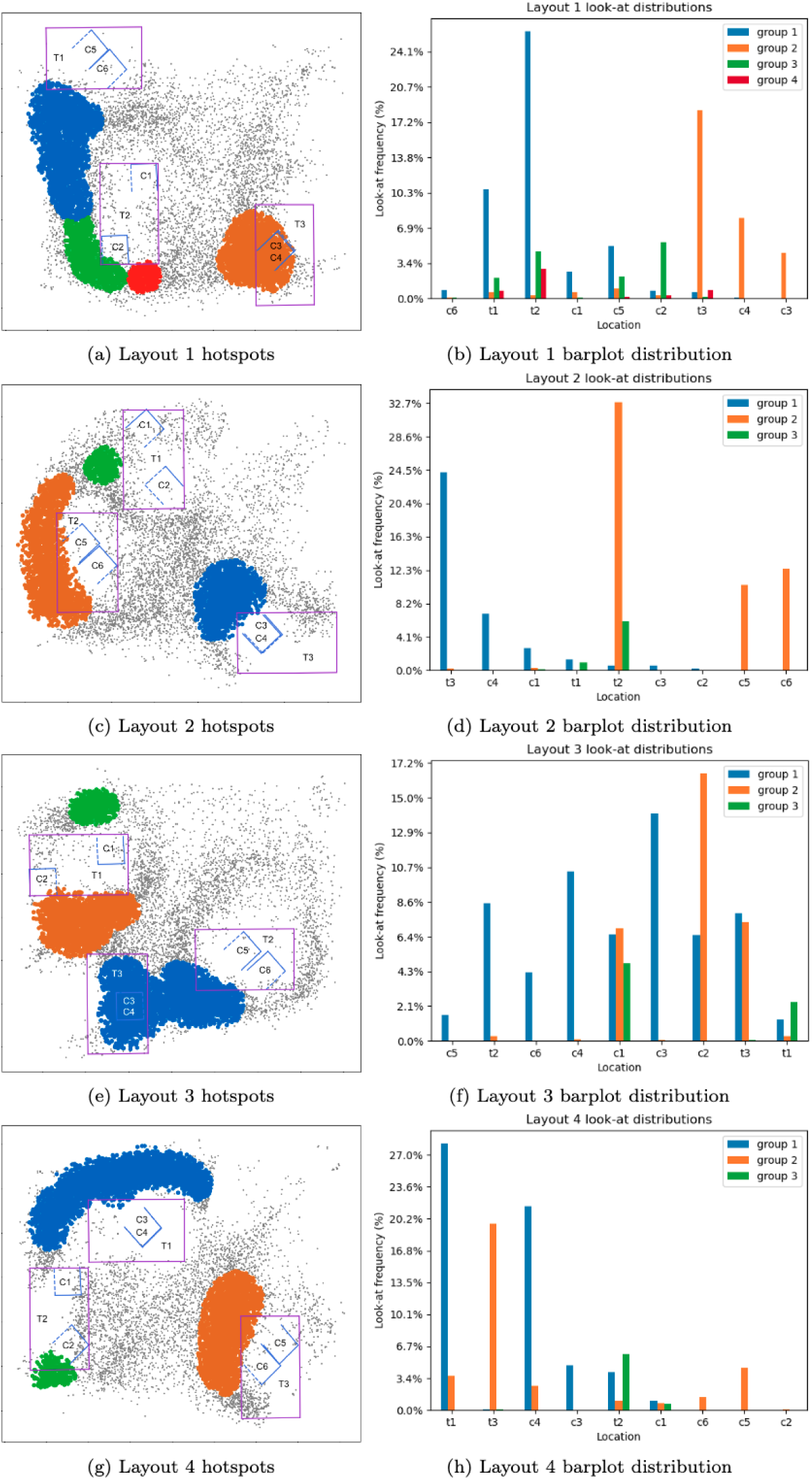
Hotspots with subject head locations are plotted on the left. Each colour corresponds to a particular cluster as defined using DBSCAN, while black dots do not belong in any cluster. Purple rectangles are tables, blue outlines are cages. Solid blue lines indicate covered faces, dotted blue lines indicate uncovered faces. Sides with no line are openings of the cages. Barplots on the right show the corresponding distribution of fixations looking at each location (table or cage) from each cluster. For example, in Layout 2, there is a table T2 with a two cages side by sie (C5, C6). The orange cluster is located at the opening of the cages and around T2. Correspondingly, the barplot shows orange to have the most frequent look-at fixations at T2, C5, C6. See Fig 1 for a clearer depiction of tables, cages, and their labels.

Although this may seem intuitively obvious, the data reminds that subjects must detect such viewpoints, then navigate to them, and orient their head/eyes for appropriate viewing. From any viewpoint, starting or otherwise, there are always a number of cage views that are occluded either by a cage wall or by another cage. There is a clear navigation aspect required in this search task to move subjects to a viewpoint from which they can inspect the cages, but this first requires detecting which viewpoints would lead to a visible cage. Thus, navigating to that viewpoint involves hypothesizing a viewing position from which cage contents are visible (a visibility affordance), then planning and executing the path to it, as well as orienting the head and eyes towards the cage opening. Such a sequence of actions is not trivial and must be planned and orchestrated. The behaviour to seek unoccluded views is not apparent in other visual search experiments, nor has it previously been recorded.

#### 3.5.3 Summary

To address the final aspect of our secondary goals, whether object placement and orientation affects viewpoint selection, we found significant difference in head movements for different target orientations, demonstrating that subjects adjust their views accordingly. In particular, we can see that side-up objects induced larger pitch angles to tilt the head down when viewing the object compared to upright objects. When objects were placed lower (on tables vs. on top of cages), subjects tended to crouch more to level their views with the height of the objects.

Finally, objects placed in occluded areas cause subjects to move around the environment to find unobstructed viewpoints to perform their search. This is a type of behaviour not previously recorded in other real-world or search experiments, yet is an integral part of any real-world search task involving occlusion.

## **4** Discussion

In this experiment, we investigated the characteristics of viewpoint selection in an active visual search task conducted in a real-world environment. Most of the active vision literature involving search has thus far focused on search when the potential targets are already in view [11, 12, 40, 42]. The few that had stimuli beyond a small fixed search space include [10, 43, 47]. However, Land et al. [43] had only three subjects, poor record of where the possible stimuli were, and no head position tracking, Franchak et al. [10] had very simplistic targets that did not require close scrutiny to identify, and Kit et al. [47] performed the experiment in virtual reality, where subjects had to use a controller to move their body. Our experiment addresses these issues by including head movement tracking, recorded location of target objects, and structures used to deliberately introduce occlusion in the real-world search environment.

To address our primary (What are the characteristics of viewpoint selection during active visual search?) and secondary (How do different factors affect viewpoint selection?) goals, we highlight four characteristics that have emerged from our results.

### People perform search with high accuracy, even with no training on the environment

The lack of evidence of any strong learning effects along with the high accuracy even from the first trial suggest that subjects are easily able to apply general searching strategies in unknown environments. It is possible that the small magnitude of significant efficiency gains, found only in target-present trials, is due to a ceiling effect, or that there were only twelve trials. Target not visible trials could be seen as a more complex task, as it involves more movement before a viewpoint with the target in sight is found. This could be the reason for stronger efficiency magnitudes for these trials specifically. It is possible that with more trials, larger efficiency gains may emerge.

However, Solbach & Tsotsos [39] found no learning effects on accuracy even with 18 trials of a same-different task. Beyond that, increasing the number of trials by a significant number would be difficult, as increasing the experiment duration may introduce discomfort to the subjects.

#### People are adept at selecting the least obstructed views to search through otherwise occluded areas with objects

Subjects seem to select views where they face the openings of cages to conduct search of the objects within them. This suggests that they are able to detect and select the views that provide more information than others. In other words, a visibility affordance can be associated with possible viewpoints.

#### People orient and position their head dependent on the placement and orientation of target objects

Subjects move their head, eyes, and body differently depending on the orientation and placements of target objects. If targets are placed lower in the environment, subjects tend to crouch more. Side-up objects tended to induce subjects to tilt their head down more than upright objects. Although we found no strong evidence that subjects were performing these actions to match their view to the presented view of the targets, our results suggest that different object placements and orientations still induce different head movements, as if they were tending towards a canonical eye/head pose most suited for recognition and analysis.

#### People will use the full range of motion given to them by their eyes and head to conduct visual search

Althought it might sound obvious that one moves their eyes and head, it is important to document the extent of such motions, and our reliance on them. It would, for example, be reasonable to think that one image of a scene might suffice to determine if a target is present (as [68] showed for simple go no-go tasks). This was not observed and the large numbers of different fixations subjects use emphasizes this point that the nature of the scene matters.

Most of this range of eye and head motion is moot when subjects perform search on a computer screen or on a table. A 2D visual search task with fixed head and eye position have no degrees of freedom. 2D search where subjects are free to move their eyes have 2 degrees of freedom (xy on display screen), and 5 if they are free to move their head (adds roll, pitch, yaw). A real-world search task with fixed body and head location have 3 degrees of freedom (xyz). Finally, a real-world search task with no restrictions on eye, head, or body movement has 9 degrees of freedom (xyz for eyes, xyz for head, roll, pitch, yaw), as mentioned in Section 1.1. Few studies [10, 28, 39] have recorded data for all 9 degrees of freedom, of which only one was for a real-world search task.

The larger range of eye gaze and head movements found in our experiment is consistent with prior literature, although most of this literature has focused on navigation and exploration [10, 28, 31]. However, when comparing gaze amplitudes between our task and Franchak et al. [10]’s, we notice that our average amplitude (48°) is much larger than theirs during their search task (28.5°) as well as during walking (19.5°). We posit that the design of the layouts in our search environment with three separate tables could cause many larger saccade amplitudes, as subjects switch to search from one table to the next.

#### Limitations

It is natural to wonder how a real-world active visual search task compares to 2D search. However, several aspects of our experiment design make such a comparison difficult. In particular, the need for subjects to determine viewpoints before they can fixate and scrutinize potential target objects in our task contained many behaviours without a directly comparable component in 2D search.

Finally, although we used somewhat familiar objects in the sense that the toys were mostly common furniture objects, we deliberately disrupted possible scene grammar cues by placing these objects in nonconventional orientations or placements. Future work could use the same objects, arranged with scene grammar in mind, to see if scene grammar guides search similarly in a real-world environment.

## **5** Conclusion

We sought to understand active visual search in a real-world environment, especially with regard to viewpoint selection when the search space is not fixed to a surface or screen in front of you. Results from this experiment bring us one step closer to understanding the characteristics of viewpoint selection during active visual search in the real world.

First and foremost, we presented a novel dataset of active visual search in the real world (https://osf.io/vqdb2/). This dataset includes target-absent trials as well as trials containing occlusions, neither of which had been recorded or documented in a real-world task before. This dataset showcased the full range of motion used by subjects from not only the eyes, but also the head, consistent with experiments in navigation and exploration in the real world. We also found little learning across trials in terms of efficiency, and no learning in terms of accuracy.

Importantly, the ability to manipulate several factors in our controlled real-world environment [54] makes it a powerful tool to facilitate our research of active vision in a more rigorous manner. By including occluding views and oriented targets, we were able to manipulate subjects’ viewpoint selection, allowing us to understand which views are more important than others to help identify target objects. Indeed, we showed that subjects are adept at finding the least occluded views to search through their environments, showing the use of a visibility affordance to select viewpoints, and would orient and move their heads depending on the placement of objects.

Such strategies uncovered from the human data further show the importance of considering patterns of behaviour using metrics like eye gaze and head movements together, rather than using only simple psychophysical metrics such as response time, accuracy, and search slope.

There are several avenues to explore from this experiment, from adding scene structure to guide search, altering the uniformity of objects distributed across locations, including retrieval as part of the search task, and so on. This experiment adds to the many that showcase the true complexities of visual behaviours executed in the real world, and emphasizes the importance of conducting vision research in more naturalistic environments to understand the inherently active component of visual tasks.

